# Faunal remains from the Upper Paleolithic site of Nahal Rahaf 2 in the southern Judean Desert, Israel

**DOI:** 10.1101/2022.05.17.492258

**Authors:** Nimrod Marom, Dariya Lokshin Gnezdilov, Roee Shafir, Omry Barzilai, Maayan Shemer

**Affiliations:** Laboratory of Archaeozoology, School of Archaeology and Maritime Cultures, University of Haifa – Haifa, Israel; Archaeological Research Department, Israel Antiquities Authority – Jerusalem, Israel; Department of Bible, Archaeology and the Ancient Near East at Ben Gurion University – Beer Sheva, Israel

**Keywords:** Upper Paleolithic, Levant, Judean Desert, Arkov-Divshon, archaeozoology

## Abstract

Nahal Rahaf 2 (NR2) is an Early Upper Paleolithic (ca. 35 kya) rock shelter in the southern Judean Desert in Israel. Two excavation seasons in 2019 and 2020 revealed a stratigraphical sequence composed of intact archaeological surfaces attributed to the ‘Arkov-Divshon’ cultural entity of the arid southern Levant. We present the faunal assemblages from the site, which are rare among the desert sites due to preservation problems. Our results suggest that the residents of the site exploited prime adult caprines (*Capra* cf. *Capra ibex*), but were also engaged in hunting of gazelles (*Gazella* cf. *Gazella gazella*), whose carcasses were selectively transported to the site and processed for within-bone nutrients. Long-range hunting trips are suggested by the body-part selection in relatively small bovids, and by the presence of species representing wetter habitat patches 20-30 km north of the site. The bi-focal emphasis on generalized gazelle and age-specific caprine hunting, is unique among Late Pleistocene sites from the Levant. The proportion of caprines increases through the stratigraphic sequence, suggesting more specialized economy through time and in inverse relations to site use intensity.

## Introduction

The Levant, as the only land bridge between Africa and Eurasia, preserves a record of faunas and hominins that have moved into it during recurrent Pleistocene expansions from both continents (e.g. Belmaker & Bar-Yosef, 2011; Goren-Inbar & Speth, 2004; Timmermann & Friedrich, 2016). The Upper Paleolithic period (50-22 ka) is known to correspond with one of those expansions, the Recent out of Africa, or the spread of modern humans into Eurasia (e.g. Douka et al. 2013; Hershkovitz et al. 2015; Boaretto et al. 2021).

The Levantine chrono-cultural sequence is quite complex and is composed of three phases: Initial, Early and Late Upper Paleolithic (Bar-Yosef and Belfer-Cohen 2010). Each phase comprises at least two geographical entities that are characterized by specific material cultural remains such as stone tool technologies, bone and antler tools, and shell ornaments. The Initial Upper Paleolithic (IUP) phase includes the Emirian, dated to as early as 50 ka at the site of Boker Tachtit (Boaretto et al. 2021), which developed into a later phase which is generally regarded as ‘IUP’ and was found, for example, at Boker Tachtit (Level 4), Ksar Akil (Layers XXV-XXI) and Üçağizli Cave (layers F-I) (Boaretto et al. 2021; Barzilai in press). The Early Upper Paleolithic (EUP) phase includes two cultural entities – the Early Ahmarian and the Levantine Aurignacian. The Ahmarian is widely distributed throughout the Levant, and seem to have northern and southern variants/facies which are reflected in the lithic industries (Gilead, 1991; Kadowaki and Nishiaki, 2016). The northern facies first appear ca. 46 kya, while the first appearance of the southern Ahmarian is dated to ca. 42 kya, and shows a clear technological development from the late Emirian. The Levantine Aurignacian, on the other hand, is dated to 39-33 kya, and considered to represent a local adaptation to of the European, Aurignacian techno-complex (Bar-Yosef, 2003; Alex et al. 2017; Marder et al. 2021; Tejero et al. 2021; Sadhir et al. 2020).

The Late Upper Paleolithic (LUP) is less well known. Culural entities that were long considered to encompass the transition from Upper-Paleolithic to Epi-Paleolithic lithic industries inclue the Atlitian, Arkov-Divshon and the Masraqan. Where the Masraqan was considered as a postdecessor of Ahmarian industries, the Atlitian and the Arkov-Divshon were associated with the Levantine Aurignacian (e.g., Gilead 1991; Marks 1981; Goring-Morris and Belfer-Cohen, 2018).

However, the new excavation at the site of Nahal Rahaf 2 revealed a preserved sequence of Arkov-Divshoon assemblages dated 39-33 ka, which attributes this cultural entity to the EUP phase (Barzilai et al. 2020; Shemer et al. forthcoming).

Subsistence practices and the paleoenvironmental context of Upper Paleolithic cultural entities has preoccupied archaeological research for many decades. However, the faunal research is incomplete and focused on particular sites, mainly large cave sites in the Mediterranean region of the Levant (Rabinovich, 2003; Stiner, 2005; Marín-Arroyo, 2013; Orbach & Yeshurun, 2019; Stiner et al., 2005; Yeshurun et al., 2021). The situation is different in the desert regions, where sites show limited stratigraphic sequences and poor preservation of non-lithic finds.

Zooarchaeological data from the Upper Paleolithic desert regions of the southern Levant are known from thirteen sites, none of which has a sample size larger than 500 identified bones (Rabinovich, 2003); some of the largest were excavated in the earlier twentieth century using recovery, retainment, and analysis methods typical of that time (Vaufrey, 1951). Moreover, the subsistence practices of the Upper Paleolithic industry endemic to the arid fringes of the southern Levant, the Arkov-Divshon culture, is known from a single taxonomic list of the fauna Ain Aqev (D-31) (Tchernov, 1976). The important and timely theme of human adaptation to marginal environments is therefore difficult to address just when and where human populations undergoing global expansion have first set foot in Eurasia.

The site of Nahal Rahaf 2 (NR2) in the Judean Desert, Israel, is unique in providing a stratified, single-period Upper Paleolithic sequence from what is today a hyper-arid region of the southern Levant. The site, discovered in 2019, comprises eight layers representing human and natural deposition (Barzilai et al., 2020). The presence of twisted bladelets and carinated scrapers associate it with the little-known Arkov-Divshon lithic industry. The site yielded an animal bone assemblage that provides an opportunity to study Upper Paleolithic subsistence behavior in the desert regions of the Levant; in fact, it is the only zooarchaeological assemblage from the region that has been recovered by sieving, comprises ~1,000 identified bones, and derives from tightly controlled chronostratigraphic contexts validated by radiocarbon dates.

This study presents a faunal analysis of the NR2 animal bone remains recovered during the 2019-2020 excavation seasons at the site. Our objective is twofold. Firstly, paleoenvironment: NR2 dates to MIS3, which preceded the LGM and is reconstructed as cool and wet, with low speleothem δ13C values in the Mediterranean zone, speleothem formation in karstic caves in the Negev (Vaks et al., 2006) and high lake stands (Torfstein et al., 2015). Recent isotopic analysis of broadly contemporary gazelle remains from Tor Hamar in southern Jordan suggest open, steppe environments (Naito et al., 2022). The faunal remains from NR2 can supplement these geographic and climatic observations and scale them to the language of human subsistence opportunities and the availability of habitats for different game animals in what is today a hyper-arid desert. We assume that during the Pleniglacial the southern Judean Desert would have been moister than today, but not at the level of the known sites excavated in the northern and western part of the region at sites such as Erq el-Ahmar and al-Quseir (Vaufrey, 1951).

Secondly, we are interested in the game procurement strategies of the Upper Paleolithic occupants of the site, especially with respect to taxonomic and demographic prey choice and carcass transport decisions. Was the hunting limited to a narrow spectrum of local taxa? Can we observe a broadening base of smaller game and birds, as can be seen in contemporary Manot Cave in the Galilee (Yeshurun et al., 2021)? Did the hunters show preference for the transportation of specific skeletal elements to camp, or were carcasses transported more-or-less complete to camp? From the few published zooarchaeological assemblages from the Upper Paleolithic non-selective hunting of gazelles, represented by complete carcasses, appears to have been the main pillar of subsistence (Stiner, 2005; Yeshurun, 2021). Is this also the rule far inside what is, today, a desert? We lack knowledge on skeletal element representation and prey demography in these regions. The faunal assemblage from NR2 provides, therefore, a unique opportunity to study human arid land subsistence in the Upper Paleolithic Levant.

### Site and Setting

NR2 is a rockshelter in the base of the steep cliffs of the Rahaf canyon, the southern Judean Desert (Fig. 1). Current environmental conditions are arid, with a mean annual rainfall between 130 and 50 mm yr^-1^. Most of the precipitation falls in short rain storms during the spring or autumn (Zituni et al., 2021). The rock shelter is a partly open space (~ 35 m^2^), at the entrance of a cave comprising two chambers that are filled with sediments and have not yet been explored. Two working seasons resulted in 9 m^2^ of excavation at the site, which included the excavation of a test pit through ~1.5 meter accumulation above bedrock. The sequence was divided into eight layers. Layers 5–8 contained Upper Paleolithic archaeological finds (Barzilai et al., 2020), dated by radiocarbon to ~39–32 ky Cal BP. Younger ages were obtained by Lazagabaster et al. (2021: Table S3), through the radiocarbon dating of the bioapatite component in two leopard phalanges from Layer 5 (30247±70; 33065±70).

**Figure 1:**
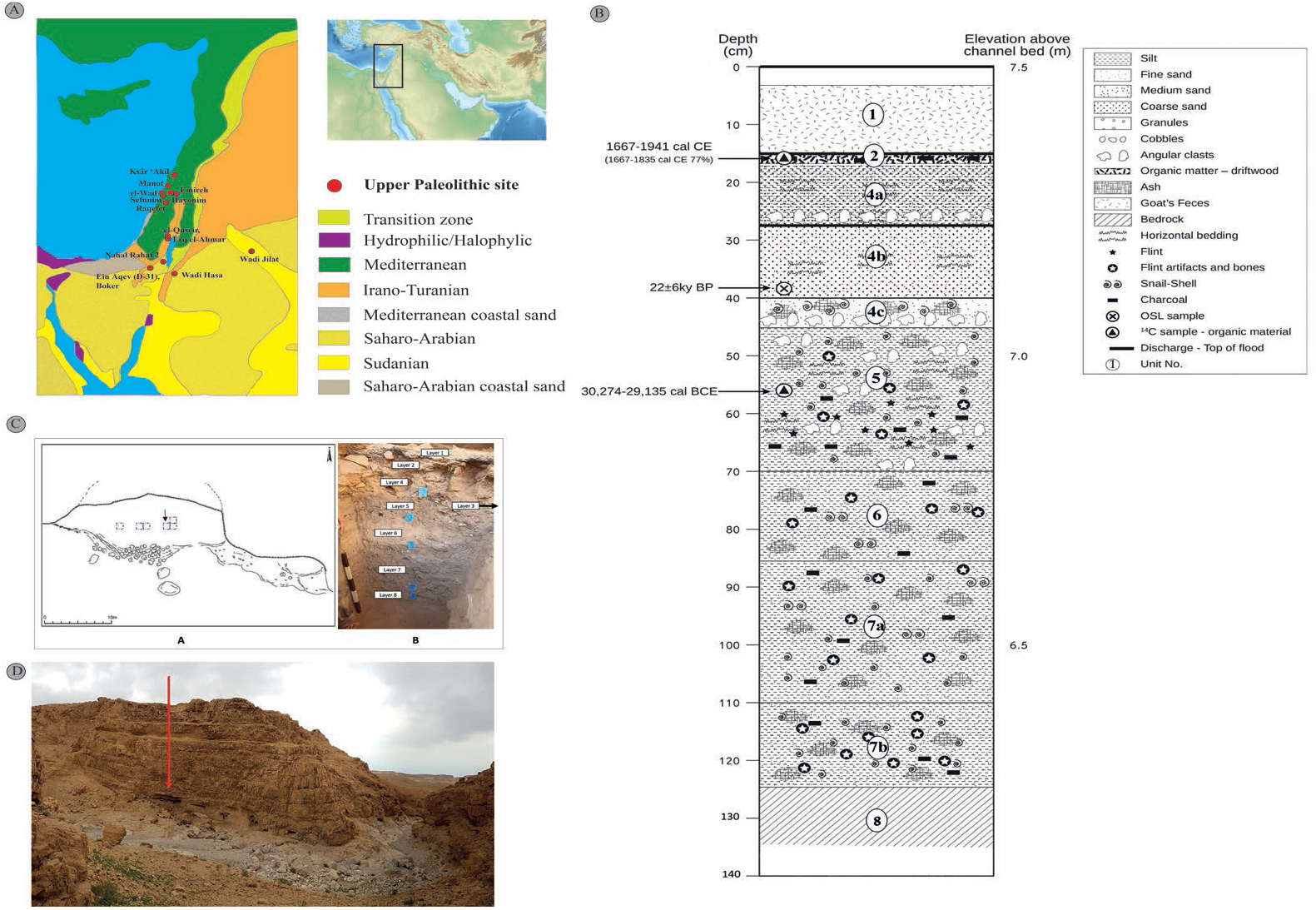
(A) Major Upper Paleolithic sites in the southern Levant. Regional map with modern phytogeographical zones and key Upper Paleolithic sites in relation to the chronology. Map of the Middle East modified from https://commons.wikimedia.org/wiki/File:Middle_East_topographic_map.png under Creative Commons Attribution-Share Alike 4.0 International license; phytogeographical map drawn by NM based on Zohary (1973). (B) stratigraphic section of NR2 sounding (Modified, after Zituni et al., 2021); (C) sketch of the NR2 rockshelter and excavation areas; photograph of sounding; (D) the rockshelter, view from the south.

Layer 8 comprises eroded bedrock with hardly any archaeological finds. These few finds, represent materials from the layers above it rather than *in situ* accumulation. Layer 7 is divided into two sub-layers, 7a and 7b based on the presence of a hearth, but in this paper is not maintained, to keep workable sample size. Combustion features, such as ashy striations and hearths were identified also in Layer 6. Archaeological layers 5—7 are extremely rich in lithic artifacts, faunal remains, mollusk shells (including perforated marine shells), and macro botanical remains (charcoal). Ochre staining occurs on both shells and artifacts. Later layers contain less evidence for intensive human activity. Small quantities of flint artifacts associated with the Arkov-Divshon industry and faunal remains were also found in Layers 4-1. The lithic artifacts found in those layers bear technological resemblance to the Arkov-Divshon lithic industries defined in Layers 5-7. Therefore, the presence of archaeological finds in the upper layers was mostly ascribed to bioturbation and post depositional erosion, although ephemeral in situ activity cannot be discounted.

Below is a short sedimentological description of the layers at the site:

<u>Layer 8</u> was exposed at the bottom of the sequence, over a very small area. It consists of a yellowish, sandy sediment ~0.1 m thick covering stone – probably bedrock. It is almost devoid of archaeological finds.

<u>Layer 7</u> loose, gray sandy sediment, ~0.4 m thick, with combustion features (a hearth) and a high density of flint and faunal remains. It is divided into two sublayers (ignored in this study) by a possible living surface.

<u>Layer 6</u> light gray/whitish sediments, ~0.4 m thick. Find density is high, but somewhat lower than the overlying Layer 5 or underlying Layer 7. A combustion feature and a living floor were found.

<u>Layer 5</u> dark gray sediment, ~0.25 m thick. Very high density of finds.

<u>Layer 4</u> coarse-to-fine grained yellowish sand, deposited in multiple flood events. Its thickness increases from north (~0.15 m) to south (~0.6 m), where it cuts the archaeological layers.

<u>Layer 3</u> pits in the north-eastern part of the rockshelter, cutting from Layer 2 (?) to Layers 4 and 5, to a maximum depth of 35 cm.

<u>Layer 2</u> burnt organic material and woody remains (~0.15 m) overlying Layer 4. Among the finds are several pottery shards and hearths. The layer was radiocarbon datedby Zituni et al. (2021) to 1667–1941 Cal CE.

<u>Layer 1</u> goat/sheep dung (~0.15 m) representing recent herding activities.

## Materials and Methods

The faunal remains from the site were collected by dry sieving all the sediments through a 2-mm aperture net during the excavation, which was carried out in 0.5 sq m grid. The bones were not washed or treated with chemicals; those that required thorough cleaning were sonicated. Specimens were identified using the comparative osteological collection of the Laboratory of Archaeozoology at the University of Haifa. When identification to species was not possible, as in the case of most shaft fragments, the specimens were assigned to one of four size classes: hare/dog, gazelle/ibex, fallow deer or aurochs-sized animals. Bones that could be thus identified to skeletal element and size-class (henceforth NISP, Number of Identified Specimens) were recorded with reference to the presence of scan sites (Lyman, 1994, fig. 7.4). Minimum number of elements (MNE) was calculated as the most frequent scan site tallied for each element. The MNE values were normalized by the number of elements in an artiodactyl skeleton to derive Minimum Number of Animal Units (MAUs). Linear measurements were taken following von den Driesch (1976). The state of epiphyseal fusion and tooth eruption and wear for caprines and gazelles were also recorded (Haber & Dayan, 2004; Munro et al., 2009; Zeder, 2006). Other parameters such as fragment length and color were noted and appear in the database (Supplement S1).

Although taphonomic analysis is not at the focus of this study, we present here basic data on bone surface modifications and bone preservation as a background check to the assumption that the assemblage reflects, by and large, human subsistence activities rather than biogenic cave deposition that is more common in the Judean Desert region (Horwitz et al., 2002; Lazagabaster, Égüez, et al., 2021). All specimens were observed with a Dinolite™ microscope using at least X5 magnification to scan for bone surface modifications such as carnivore damage, cut marks, and percussion marks. Burning, weathering (Behrensmeyer, 1978) and breakage morphology (Villa & Mahieu, 1991) were also recorded.

Statistical tests were computed in PAST 4.09 (Hammer et al., 2001), unless noted otherwise. Rarefaction diversity tests utilized library ‘rareNMtests’ (Cayuela & Gotelli, 2014), and components of the ‘tidyverse’ library were used for data visualization (Wickham et al., 2019). The calculations of NISP per element, counts of diagnostic zones, and MNE per element for each layer were automated using a short code in R (R Core Team, 2021), to facilitate replicability and eliminate errors. The code and data are available as an R library (‘SEAcalc’, https://github.com/nmar79/SEAcalc).

## Results

### Assemblage formation

The faunal assemblage from NR2 comprises 987 bones identified to skeletal element and at least size class taxon. Bone surface modifications in NR2 are almost entirely anthropogenic, and include cut marks (N=39), percussion marks (N=3), and burning (N=137) (Table 1). Non-human bone surface modifications are extremely rare, comprising three cases of carnivore and one of rodent gnawing. Bone weathering levels above 2, which suggest prolonged surface exposure before burial, is common in Layers 2–4. Layers 5—7, which include intact archaeological contexts, are almost free of indications for mechanical diagenetic processes such as dry fracturing and weathering, or biogenic processes such as gnawing. Bone preservation at the site can be quantified by correlating CT derived bone mineral density, (Lam et al., 1999: table 1, values for *Rangifer tarandus*), and the MAU calculated from each scan site location. The results (Spearman’s r = 0.21, p = 0.07) suggest a weak and not statistically significant effect of density values on the representation of diagnostic regions in the bones.

**Table 1:** Bone surface modifications and fracture morphology at NR2. Percentages were calculated out of the NISP, except for the fracture morphology where percentages were calculated out of the sum of observed fracture morphologies.

| Data | Layer 2 | Layer 3 | Layer 4 | Layer 5 | Layer 6 | Layer 7 |
| --- | --- | --- | --- | --- | --- | --- |
| NISP | 39 | 82 | 60 | 240 | 99 | 465 |
| Cut | 1 (2%) | 2 (2%) | 2 (3%) | 10 (4%) | 4 (4%) | 20 (4%) |
| Percussion |  |  |  |  |  | 3 (0.6%) |
| Burning |  |  |  |  |  |  |
| Charred | 2 (5%) | 6 (7%) | 7 (12%) | 38 (16%) | 29 (29%) | 43 (9%) |
| Calcined |  | 1 (1%) | 4 (7%) | 16 (7%) | 5 (5%) | 16 (3%) |
| Gnawing |  |  |  |  |  |  |
| Rodent |  |  |  |  |  | 1 (0.2%) |
| Carnivore | 1 (2%) |  |  | 1 (0.4%) |  | 1 (0.2%) |
| Weathering |  |  |  |  |  |  |
| Stage 2 | 4 (10%) | 17 (20%) | 9 (15%) |  |  | 3 (0.6%) |
| Fracture |  |  |  |  |  |  |
| Dry, <50% |  | 2 (9%) | 1 (3%) |  |  | 2 (1%) |
| Dry, >50% | 1 (20%) |  | 2 (7%) | 13 (12%) | 7 (11%) | 16 (9%) |
| Green, <50% | 3 (60%) | 7 (33%) | 9 (33%) | 1 (1%) |  | 5 (2%) |
| Green, >50% |  | 3 (14%) | 10 (37%) | 61 (58%) | 39 (61%) | 81 (47%) |
| Mixed, <50% | 1 (20%) | 5 (23%) | 3 (11%) |  |  | 10 (5%) |
| Mixed, >50% |  | 4 (19%) | 2 (7%) | 29 (27%) | 17 (26%) | 58 (33%) |

### Taxonomic composition

The faunal remains from NR2 comprise 987 bones identified to size (NISP = 524) or biological (NISP = 463) taxon. The two dominant taxa are goat (probably the Nubian Ibex, *Capra* cf. *Capra ibex,* NISP = 201, 20%) and gazelle (*Gazella* sp., NISP = 159, 16%). It is difficult to identify the gazelles to the species level: today, both Dorcas gazelles (*Gazella dorcas)* and Mountain gazelles (*Gazella gazella)* are extant in the region (Yom-Tov & Mendelssohn, 1975). It was suggested by Tchernov that the Dorcas gazelle has expanded to the southern Levant only in the Holocene (Tchernov et al., 1986/7), but this has not been demonstrated conclusively. Horwitz and Goring-Morris (2000), grappling with the same taxonomic issue in their publication of the Epipaleolithic fauna from Upper Besor 6, have compared measurements of recent Dorcas and Mountain gazelles from Israel and have found a substantial overlap that does not support taxonomic identification to species based on linear measurements of post-cranial bones. They suggest, however, that horncore morphology and base length and breadth measurements are more effective in distinguishing these gazelle species. A near complete horncore from NR2 (NR-0321; L114, B1147, Layer 7) has a round cross section (Fig. 2) and appears to conform both morphologically and metrically (Fig. 3) with a female Mountain gazelle (*Gazella gazella).*

**Figure 2:**
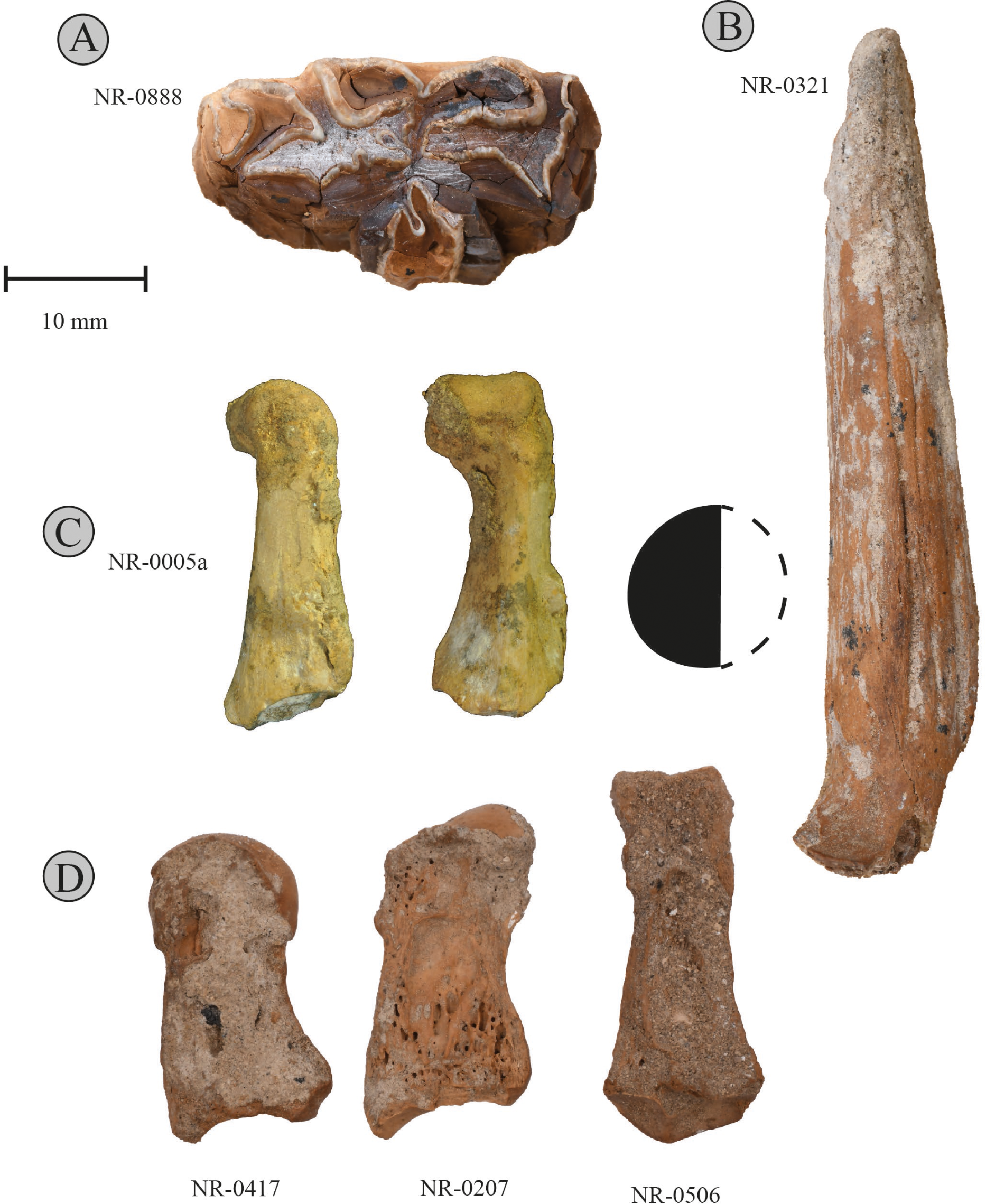
(A) right-sided mandibular premolar of an assinine equid (Layer 7a); (B) female *Gazella gazella* horncore (Layer 7), with cross section of the posterior part of the base; (C) leopard (*Panthera pardus)* phalanges (Layer 5). (D) longitudinally-split phalanges. Photographs by Roee Shafir.

**Figure 3:**
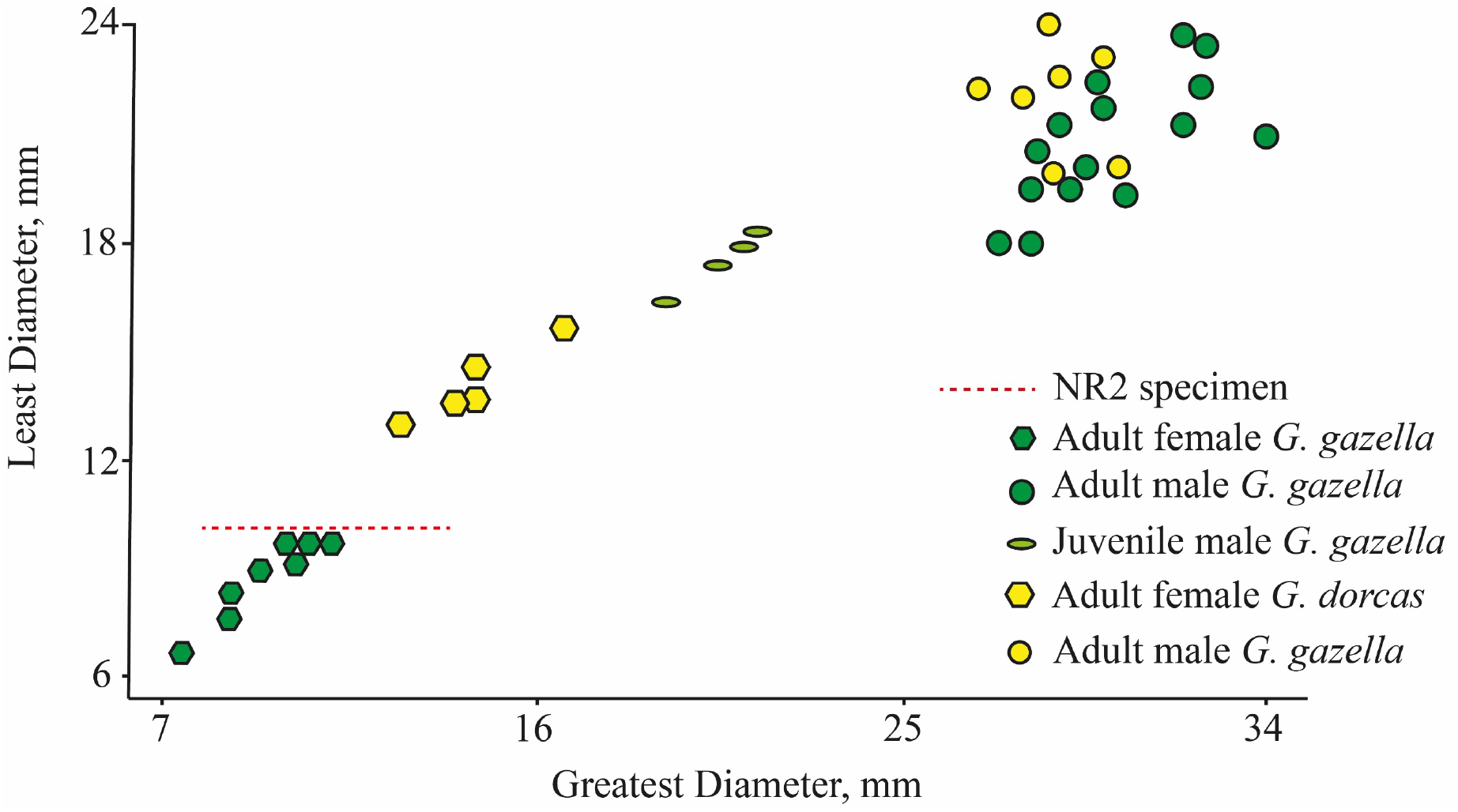
Gazelle horn core dimensions. Modified from Horwitz & Goring-Morris (2000: fig. 5).

Other ungulates are represented by few remains. A small equid is represented by seven specimens, five of which are from Layer 7. The only potentially diagnostic equid specimen is a right p3/p4 (NR-0888, Layer 7a; Fig. 2). The tooth is not well-preserved, but a V-shaped lingual fold and small dimensions (L = 29.8 mm) suggest that it belonged to an asinine equid. Unfortunately, enamel fold morphology of the occlusal surface is not diagnostic in premolars, and so the specimen could have belonged to a hemione (*Equus hemionus*), a European wild ass (*Equus hydruntinus)* or an African wild ass (*Equus africanus).* The first two species are very close genetically, and their dental morphology is extremely variable (Orlando 2019). Two of the equid specimens, an atlas (NR-0327, Layer 5) and a proximal metapodial fragment (NR-0709, Layer 7b), bear cut marks. Fallow deer (*Dama mesopotamica,* NISP = 2) is represented by an unfused calcaneus (NR-0991, Layer 4), bearing a cut mark, and a proximal radius fragment (NR-0201, Layer 7a). Aurochs, *Bos primigenius,* presence is attested by a fragment of a cervical vertebra (NR-0545, Layer 5) and a carnivore scored distal tibial fragment (NR-0999, Layer 7a). A broken m3 (NR-0060) and a proximal tibial shaft (NR-0153), both from Layer 3, belonged to a large antelope, which, unfortunately, cannot be identified to genus.

Carnivorans are represented by two leopard (*Panthera pardus)* phalanges (NR-0005a, NR-0006a, L109, Layer 5), and by the fragment of a *Canis* sp. cervical vertebra (NR-0230, Layer 7b). Other mammalian groups include lagomorphs, represented by the Cape hare (*Lepus capensis,* NISP=22, 2.2%), and rodents (Rodentia, NISP = 38). The latter are most frequent in the archaeological Layers 5 and 7 and are represented largely by postcranial bones that could not be securely identified to taxon; presence of *Gerbillus, Microtus,* and *Sekeetamys* is carefully suggested. Bird remains (NISP = 22, 2.2%) are fairly diverse, and include rock pigeons (*Columba livia,* NISP = 2; Layer 3), chukar partridges (*Alectoris chukar,* NISP = 4, Layers 2, 4, 5), and single bones of goose (*Anser* cf. *Anser albifrons,* coracoid, NR-0075, Layer 3), a corvid (Corvidae gen. et sp. indet., tibiotarsus, NR-0547, Layer 7b), an Eurasian coot (*Fulica* cf. *Fulica atra,* coracoid, NR-0208, Layer 7b), corncrake (*Crex* cf. *Crex crex,* tibiotarsus, NR-0137, Layer 2), and Eurasian griffon vulture (*Gyps fulvus,* phalanx 3, NR-0184, Layer 7a). Finally, two reptile bones identified as a colubrid snake and as an agamid lizard have been recorded in Layer 7.

### Diversity

A comparison of taxonomic diversity patterns was carried out using an individual rarefaction analysis on the taxonomic frequencies of the three lower strata, using the Hill number values at NISP = 33 (the smallest NISP from among the archaeological deposits, Layer 4; Layers 2 and 8 were omitted because of their small sample size), with q = {1,2,3}(Chao, Chiu, et al., 2014; Chao, Gotelli, et al., 2014; Jost, 2019) (Table 3, Fig. 3). The results suggest virtually similar diversity in terms of both evenness and richness between Layer 5 and Layer 7. Layer 6 is much less diverse. Although taxonomically more diverse assemblages characterize Layers 5 and 7 in relation to Layer 6 when sample size is accounted for by rarefaction, we suspect this has to do with the relatively smaller excavation area of the latter. The excavated remains represent a single activity context around a hearth, which may not be on par with the more heterogeneous contexts excavated in Layers 5 and 7. The diversity of Layers 4 and 3 is not less than that of those preceding them, and that bear more evidence for intensive human occupation.

**Table 2:**
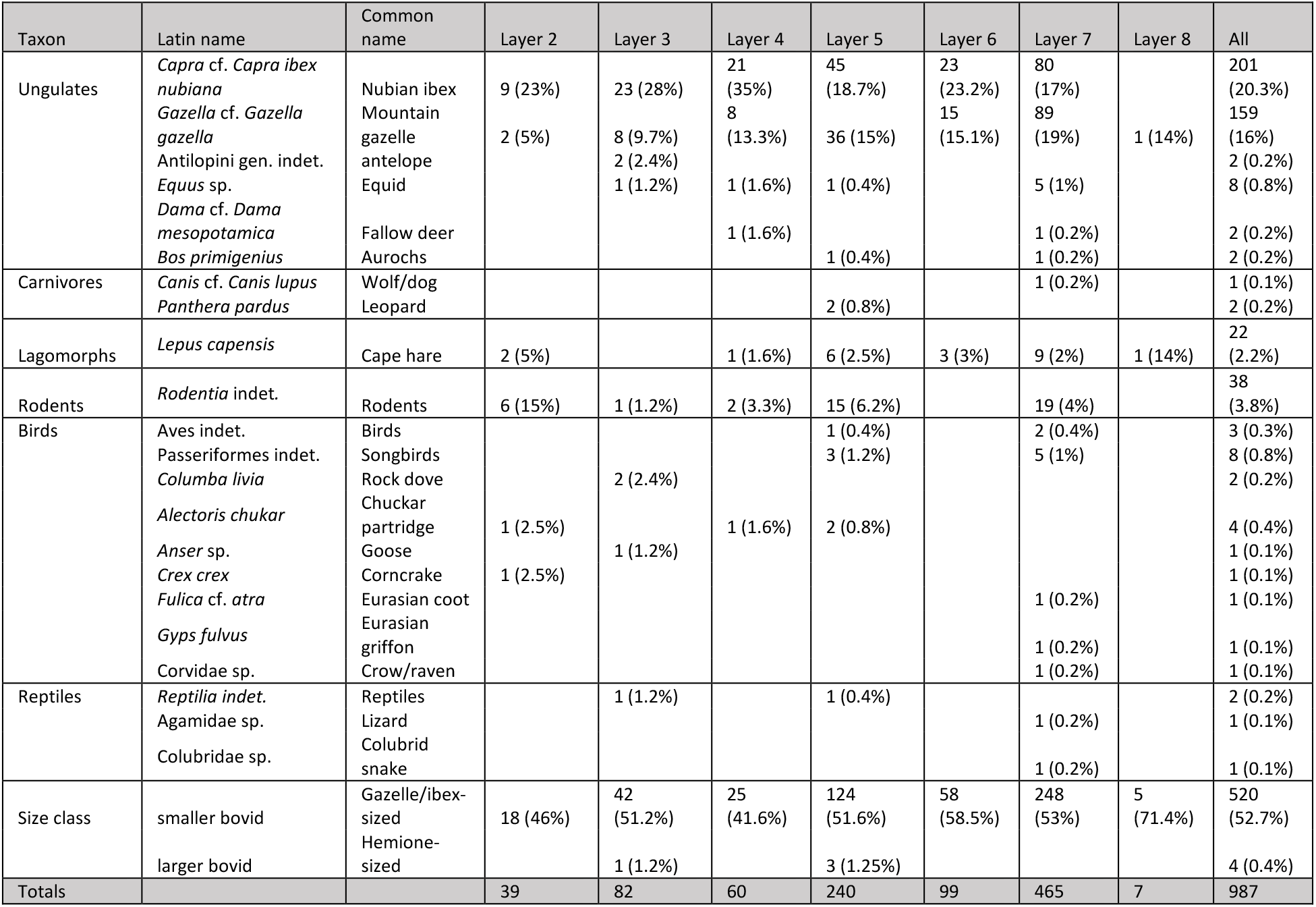
NISP counts and percentages.

**Table 3:** Hill numbers reflect the ENS (“effective number of species”) under different values of q, with q=0 reflecting richness, q=1 the exponent of Shannon diversity, and q=2 the reciprocal of Gini-Simpson index. The values were calculated using individual rarefaction for NISP=33, the size of the smallest sub-sample (Layer 4).

| Context | q = 0 | q = 1 | q = 2 | Capra : Gazella |
| --- | --- | --- | --- | --- |
| Layer 3 (N=37) | 4.77 | 2.55 | 1.96 | 2.87 |
| Layer 4 (N=33) | 6 | 2.87 | 2.13 | 2.62 |
| Layer 5 (N=97) | 4.62 | 2.91 | 2.46 | 1.25 |
| Layer 6 (N=42) | 2.99 | 2.38 | 2.16 | 1.53 |
| Layer 7 (N=198) | 4.74 | 2.86 | 2.41 | 0.89 |

Beyond general comparisons of diversity patterns, focus on the two major ungulate prey taxa, gazelle and ibex, shows change along the stratigraphic sequence. The odds shift along the stratigraphic sequence from 0.89 in Layer 7, to consecutive 1.53, 1.25, 2.62, 2.87, and 4.5 for Layers 6 to 2 in a significant linear trend (Cochran-Armitage Test for Trend run with library ‘DescTools’ (Andri 2021; Layers 2-7: Z = −2.61, p-value = 0.009)(Table 3; Fig. 5). A comparison of the number of gazelles and caprines in the archaeological Layers 7-5 and Layers 4-2 yields a statistically significant difference (Fisher’s exact test p = 0.0004) with moderate-weak effect size (Contingency C=0.18). This suggests that higher layers, which show no evidence for intensive occupation, also include relatively more caprine remains.

**Figure 4:**
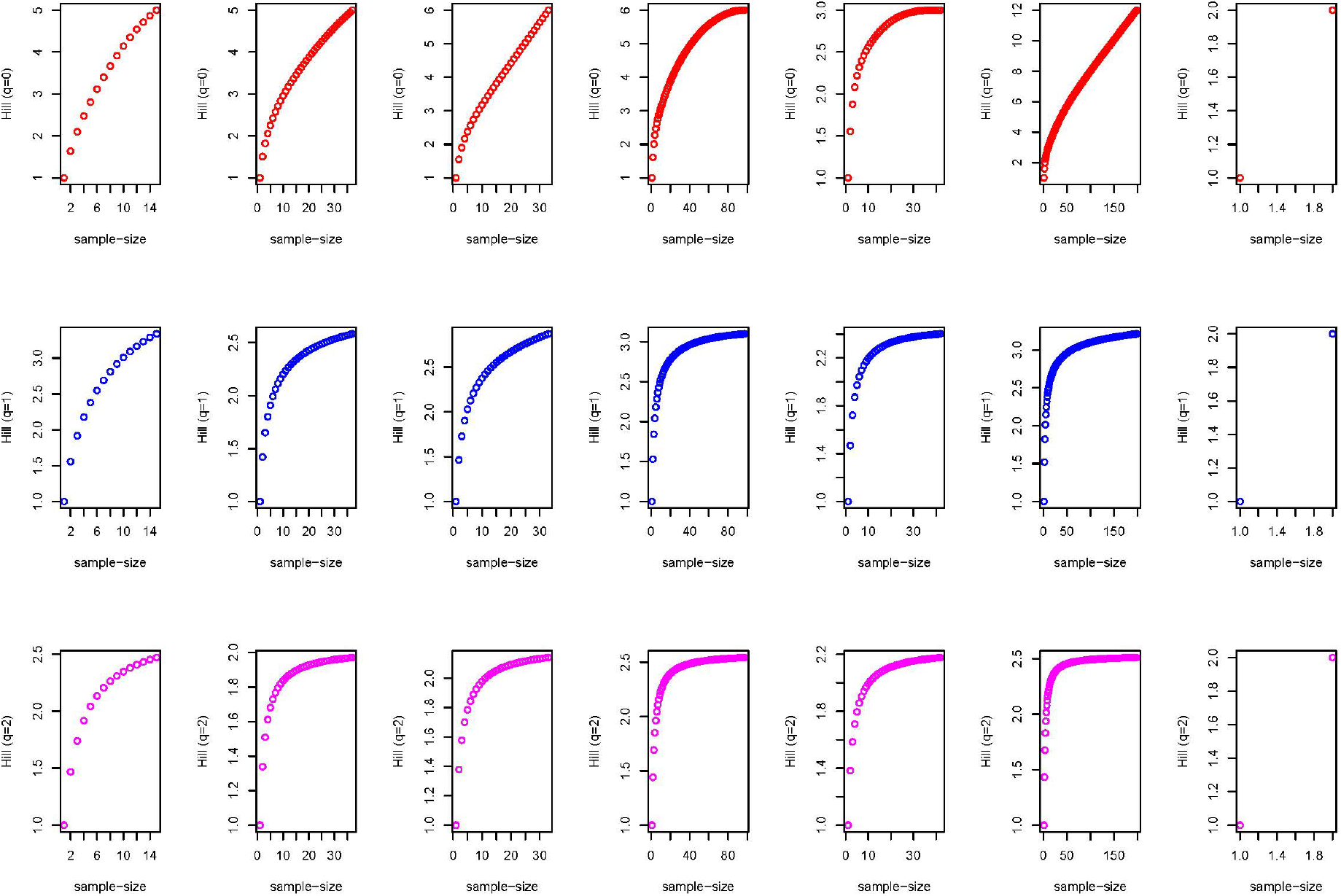
Individual rarefaction curves for Layers 2-8 (columns) for q = 1,2,3 (rows). X-axis represents the sample size in NISP, y-axis are Hill numbers.

**Figure 5:**
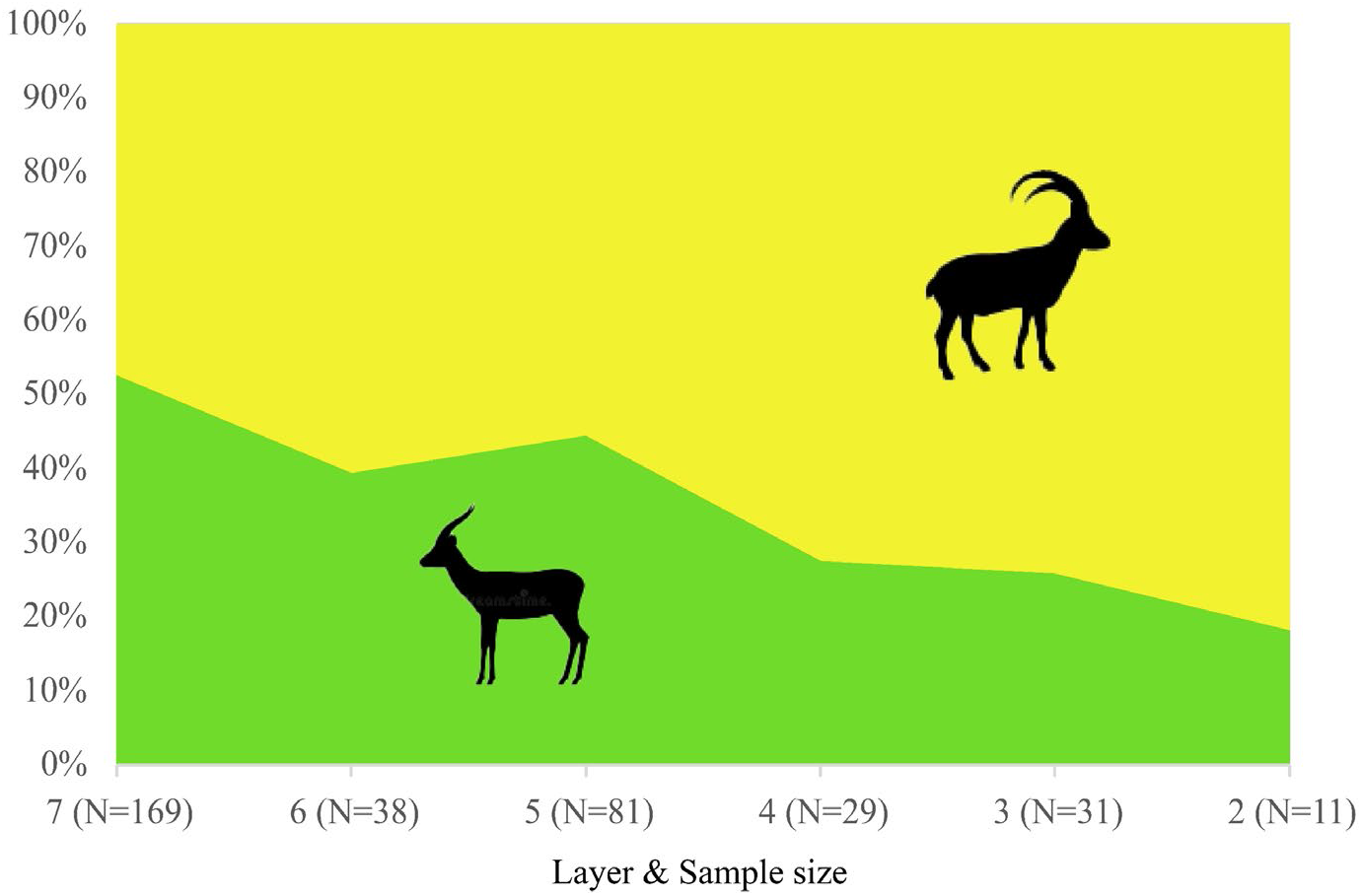
The changing ratio of caprine to gazelle bones throughout the sequence in NR2 constitutes a statistically-significant trend (Cochran-Armitage Test Z = −2.61, p-value = 0.009).

### Age-at-death and sex ratios

Age-at-death could only be estimated for gazelle and caprine epiphyses (Tables 4–5) and teeth (Table 6) when pooled across all layers. For caprines, epiphyseal fusion data suggest a culling peak at fusion stages C and, especially, D, which are young and prime adults between one and 2.5 years of age (Zeder, 2006). Tooth eruption and wear observations (N = 8) likewise suggest that individuals who had died during the second to the fourth year of their lives: There are no deciduous teeth and only one P4 is unworn. The data for gazelles are too sparse, but adults (fused bones and permanent dentition) constitute the greater majority of the ageable sample. In the case of phalanges, five out of 17 (29%) represent animals that died as fawns. This is similar to data observed in recent herds (Baharav, 1983) and in the early Upper Paleolithic assemblage in Manot, Upper Galilee (Yeshurun et al., 2021).

**Table 4:**
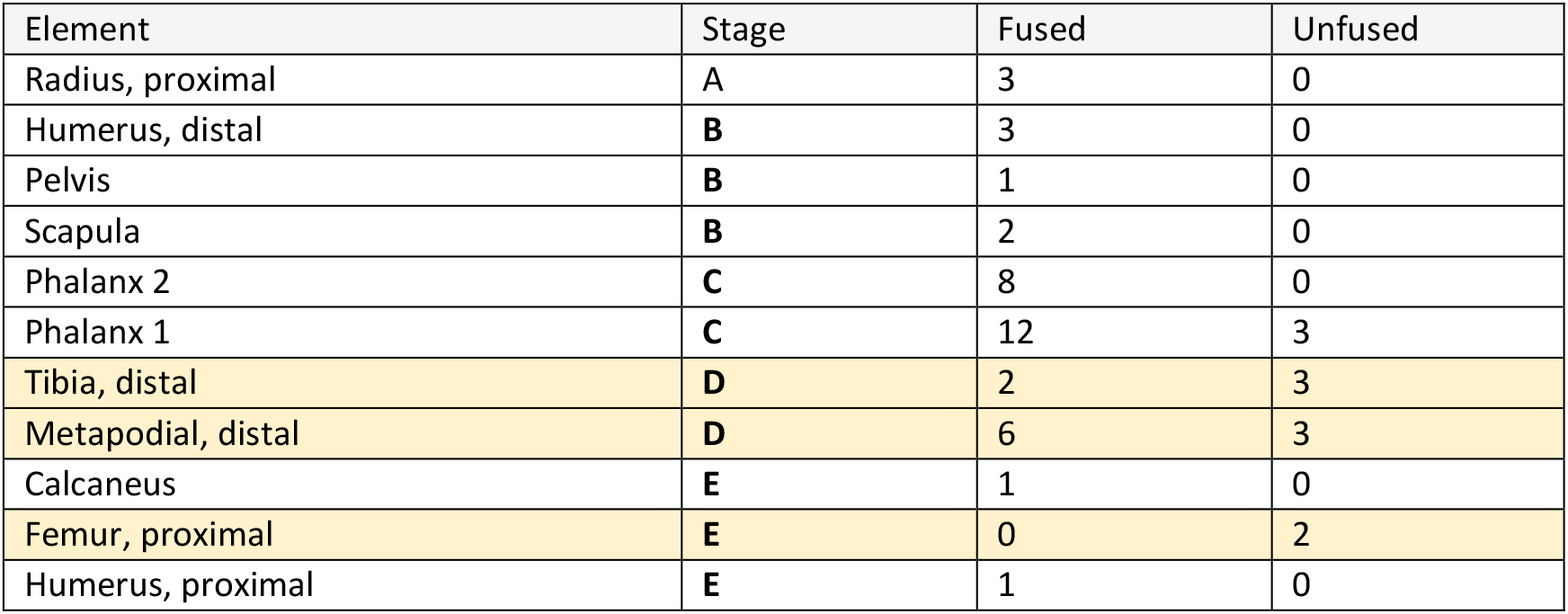
Fused and unfused bones of caprines and gazelles for all layers. Fusion stages are for goats (Zeder, 2006).

**Table 5:** Epiphyseal fusion data for gazelles, grouping after Munro et al. (2009)

| Element | STAGE | Fused | Unfused |
| --- | --- | --- | --- |
| RADIUS, PROXIMAL | II | 3 | 0 |
| HUMERUS, DISTAL | III | 1 | 0 |
| PELVIS | III | 2 | 0 |
| SCAPULA | III | 1 | 0 |
| PHALANX 2 | III | 3 | 4 |
| PHALANX 1 | III | 14 | 1 |
| TIBIA, DISTAL | V | 1 | 0 |
| METAPODIAL, DISTAL | VI | 2 | 0 |
| FEMUR, PROXIMAL | VI | 1 | 0 |

**Table 6:** Tooth wear stages (Haber & Dayan, 2004) for caprines and gazelles.

| Capra | Tooth | TWS | Age category |
| --- | --- | --- | --- |
| NR-1028 | p4 | 1 | young adult |
| NR-0263 | m3 | 2 | young adult |
| NR-0647 | p4 | 2 | young adult |
| NR-0812 | m3 | 3 | prime |
| NR-0172 | m3 | 4 | prime |
| NR-1044 | p2..4 | 4 | prime |
| NR-0048 | m3 | 3 | prime |
| NR-0011 | m3 | 5 | old |
| Gazella |  |  |  |
| NR-1066 | dp3 | 1 | juvenile |
| NR-0216 | M3 | 3 | prime |

Sex ratios for either gazelles or caprines are obscured by very small sample sizes. For gazelles, the sex ratios appear even or slightly biased towards females, based on pubis bone morphology (Table 7); to this we can add the single female horn core discussed above. The distribution of distal first phalanx breadths (Fig. 6; for measurements see S2) shows bimodality that may likewise hint at a higher female representation. Among caprines, where no horn cores and only a single pubic fragment were found, we have just clues from the size distribution of the distal first phalanx breadth measurements for a male biased sex ratio. We cannot overemphasize that this evidence does not constitute solid evidence due to sample size limitations and the weak sexual dimorphism of the phalanges.

**Table 7:** Sex frequencies by pubis bone morphology.

| Taxon\Sex | Male | Female |
| --- | --- | --- |
| Capra |  | 1 |
| Gazella | 3 | 4 |

**Figure 6:**
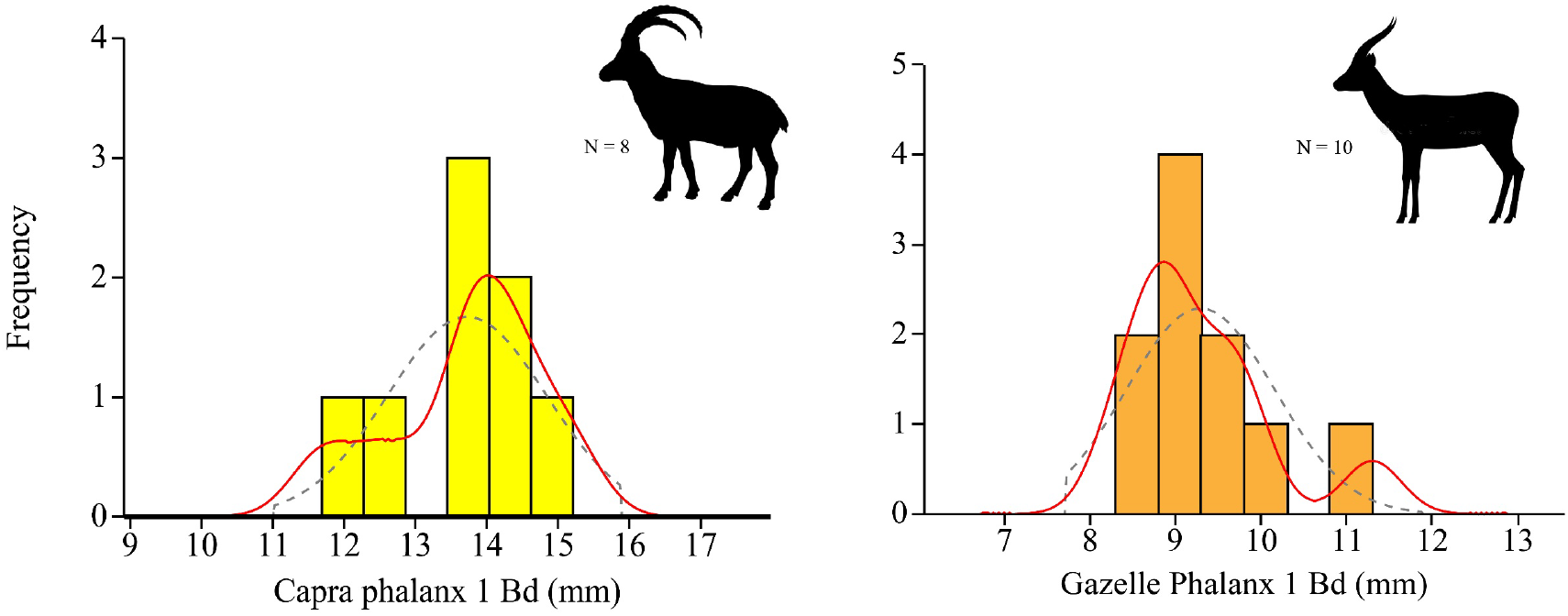
First phalanx distal breadth measurement distribution for caprines and gazelles, all layers.

### Skeletal element representation and butchery

Skeletal element analysis could only be performed for gazelles, ibex and similarly-sized bovids at NR2. When only elements identified as to biological taxa are compared, the resulting skeletal element profile is identical, and is head and foot dominated (MAU = 4,1,2,3,4 | head, axis, forelimb, hindlimb, feet). This result would be ascribed to a bias introduced by density mediated attrition affecting long bone epiphyses, which are the parts in the skeleton better identified to biological taxon (e.g., Marean et al., 2004). However, the representation of different regions in the long bones of smaller bovids is not correlated with their density in a statistically significant way, and the effect is weak (Spearman’s r = 0.21, p = 0.07; the full skeletal element representation data of smaller bovids in the assemblage appears in Supplement S3). In view of the similarity in gazelle and ibex profiles, and to accommodate the identified shaft fragments, we pooled the specimens identified as to genus (*Gazella* or *Capra)* together with the bones that were identified only as smaller bovids. Context-wise, only the two archaeological layers with the largest number of identified specimens merited separate plots (Fig. 7).

**Figure 7:**
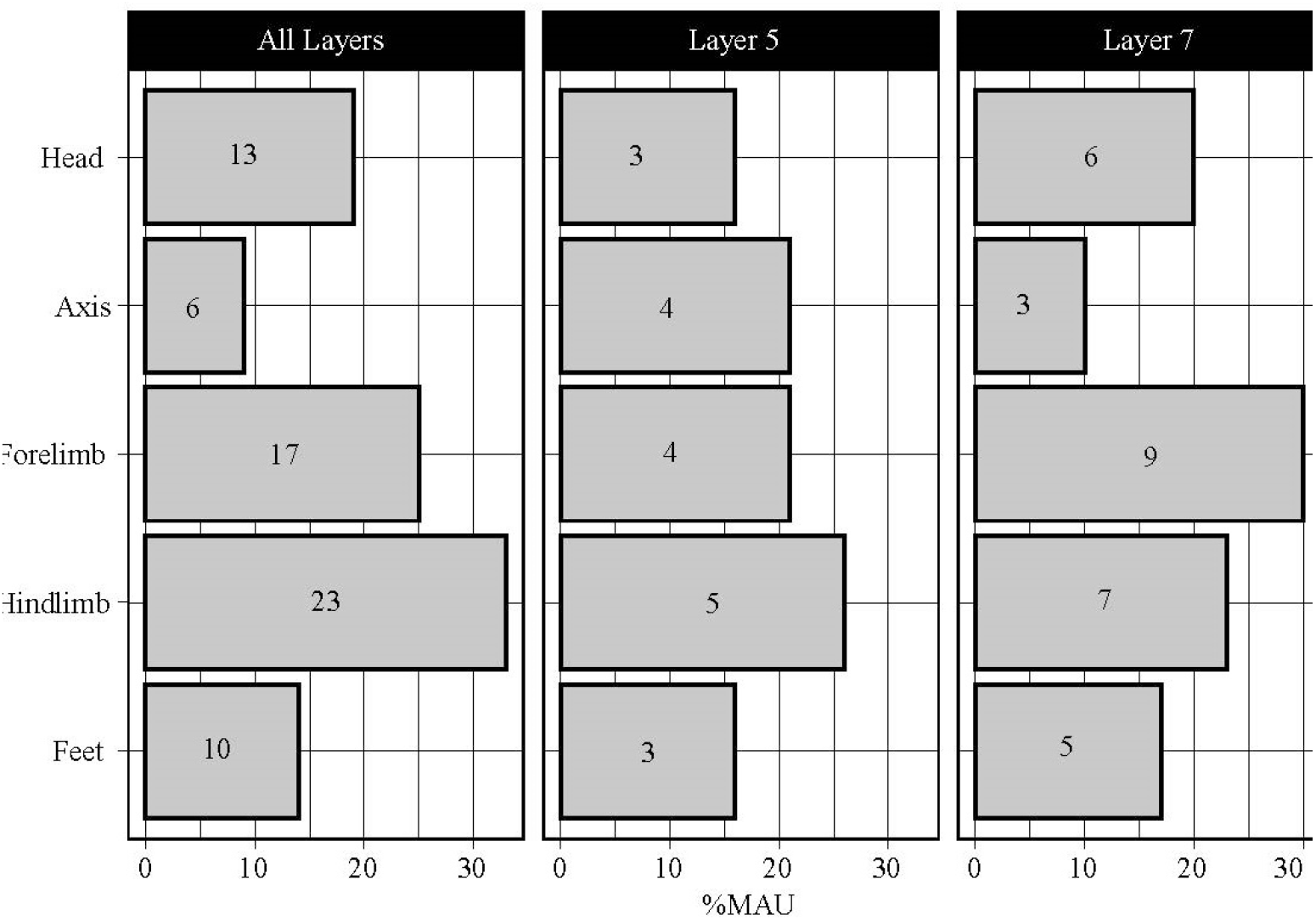
Skeletal element abundance profiles by anatomical region for caprine and gazelle sized bovids. Bar height is the percentage of the total MAU for all regions; numbers on the bars are frequencies. Head = cranium and mandible; axis = ribs and vertebrae; forelimb = scapula, humerus, radius and ulna; hindlimb = pelvis, femur, tibia, and tarsals; feet = metapodials and phalanges.

With the skeletal element abundance profile calculated for all smaller bovids and including long bone shaft fragments, the head-and-foot dominated pattern disappears. Both the synthetic profile, which pools the observations from all layers, and the specific profiles for layers 5 and 7 show a similar pattern, dominated by upper fore- and hind-limbs. For the synthetic profile, we used the highest MAU (for the hindlimbs, MAU=23) as the expected value for all skeletal regions, obtaining a statistically significant result (Chi square = 25.82, DF=4, p<0.001), which suggests that the observed skeletal element abundance profile is unlikely to represent a random sample from a faunal assemblage composed of complete skeletons. An examination of the Chi-squared standardized residuals suggests a statistically-significant (>abs(2.5)) under representation of the axis (−3.54) and feet (−2.71) regions. In summary, the skeletal element abundance profiles from archaeological layers 5 and 7, and the complete assemblage of identified bones from all layers suggest selective transport of high-utility limb bones and heads to the site. Significantly, the SEA profiles from Layers 2-4 are like those of the deeper layers that yielded richer archaeological deposits.

Bone butchery consisted of bone breakage for marrow, which we infer from the extensive fragmentation of the assemblage and the dominance of ‘green’ fractures indicating breakage of fresh bones. There is no correlation, however, between a marrow index (Jones & Metcalfe, 1988) and fragmentation intensity, as calculated from the NISP:MNE ratio (Spearman’s r = 0.29, p = 0.40); likewise, the grease index (see also Binford, 1978, pp. 23–32; see also Morin, 2007) is not correlated with spongy bone fragmentation intensity (Spearman’s r = −0.36, p = 0.33). Humans are implied as the agents of breakage because carnivore gnawing marks, digestion marks, shaft cylinders or coprolites are rare or absent from the assemblage, and few percussion marks were recorded (N=3, all in Layer 7). Notably, many (NISP = 19) of the first and second phalanges of smaller bovids were longitudinally split (Fig. 2).

## Discussion and conclusions

The faunal assemblage from NR2 represents an anthropogenic deposition, dominated by smaller bovids. Species other than ibex and gazelle are rare and include a handful of specimens identified as an asinine equid, wild cattle, fallow deer, some birds, and the carnivorans wolf and leopard. The zooarchaeological analysis suggests that Layers 7-4 all show similar patterns of taxonomic diversity and skeletal element profiles. In Layer 4, ephemeral occupations between fluvial depositions are possibly reflected by the faunal assemblage. These show a greater focus on caprine hunting, and albeit the chronology of this layer is not well secured, it is clearly later than that of Layer 5-7. We assume for simplicity that the faunal remains from Layer 3 represent disturbed archaeological remains.

Circling back to our first research question, which is about the local climatic conditions in the Pleniglacial (MIS 3), the assemblage does not support a reconstruction of significantly wetter conditions than today in the southern Judean Desert. The most common taxa are ones that occur in arid environments, including the equid, gazelle, and goat/ibex. Taxa that prefer non-arid environments, such as the fallow deer and wild cattle, are represented by a handful of specimens. Their presence suggests xeric Mediterranean habitats elsewhere in the region of the Judean Desert, a reconstruction which is congruent with the presence of African crested rat (*Lophiomys imhausi maremortum)* in the Cave of Skulls (*terminus ante quem* of ~42 kya)(Lazagabaster, Rovelli, et al., 2021); and large deer phalanges from the same site (SK-461, 42,856±457 (2σ) calBP), from Nahal Hever (HE-241, 36,665±313 (2σ) calBP), and from Nahal David (EG-034, 32,579±499 (2σ) calBP), all of them now-arid canyons some 20-30 km north of NR2. Similarly, the presence of birds such as the Eurasia coot and a goose may suggest occasional encounters near wetland patches during the spring or autumn migrations. The rarity of these taxa, however, speaks as loudly as their presence: It appears that during their stay at NR2, human groups were targeting gazelles and ibex, which can exist under current arid conditions within walking distance from the site. The rarity of taxa typical of slightly wetter environments at NR2 suggests to us that such habitats were probably limited to the northern and central Judean Desert.

Comparison of the taxonomic composition at NR2 with contemporary faunas is difficult. The few Upper Paleolithic sites from the Negev (Ain Aqev D-31, Boqer D-100, Abu Noshra I and II) and the Judean Desert (Erq el-Ahmar B—F, el Quseir C—D, Masraq a-Naj) yielded very small assemblages, or were otherwise published as taxonomic lists nearly a century ago, with no indication to overall NISP or recovery by sieving. This leaves us with a minimalistic comparison of the mean number of ungulate taxa in Upper Paleolithic assemblages from the Negev and the Judean Desert, the Mediterranean region of Israel, the Jordan Valley and Transjordan (Fig. 8, Supplement S4), which was facilitated by the detailed dataset compiled by Rabinovich (2003). A clear difference in richness appears between the northern and desert sites, and NR2 is clearly located closer to the mean number of taxa typical of the desert sites, although the number of ungulate taxa in it is higher than that mean. The dominant taxa in the assemblages from the region are ibex and gazelles, as in NR2, and unlike the northern site in which, unsurprisingly, fallow deer and wild boar constitute dominant faunal elements after gazelles.

**Figure 8:**
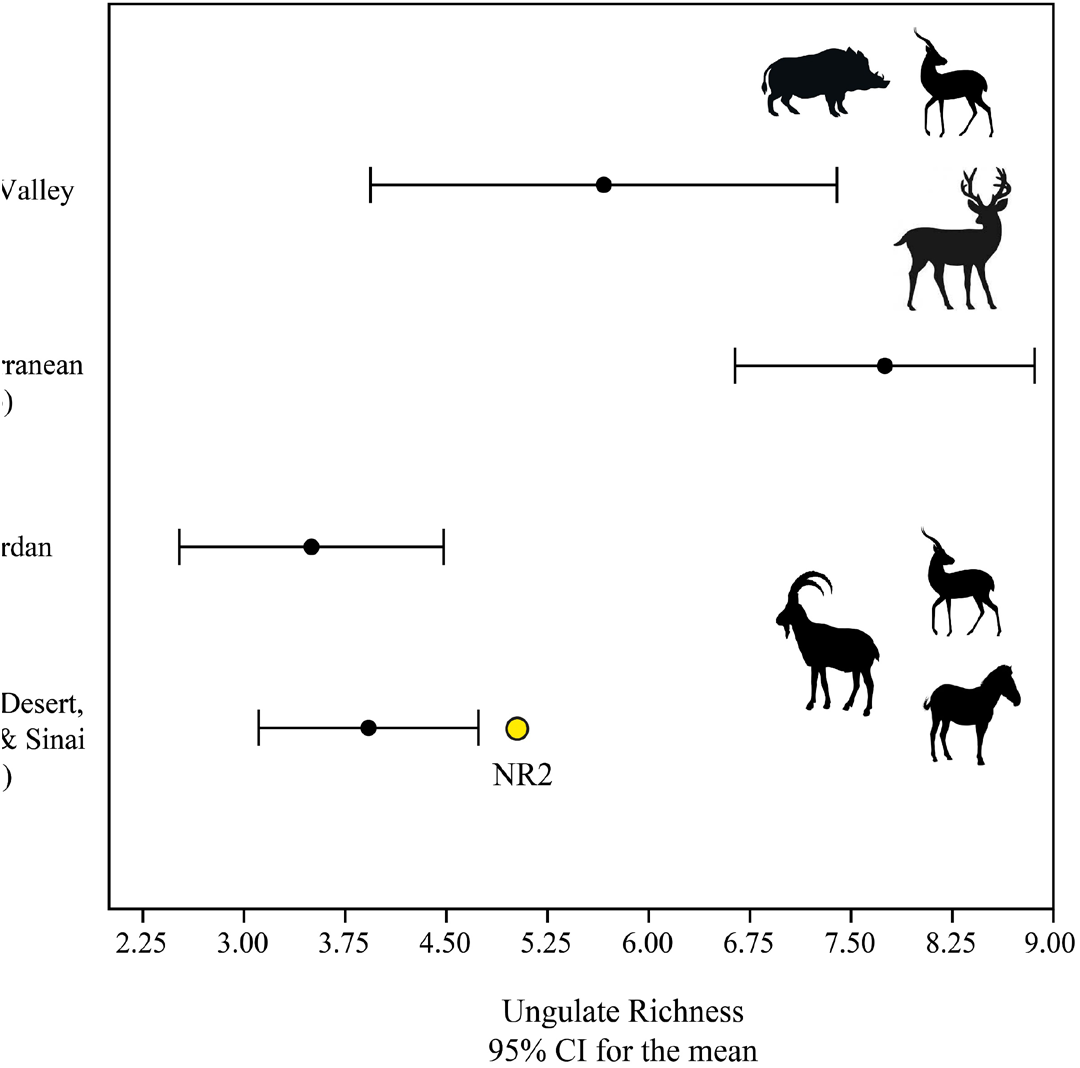
Ungulate richness with 95% CI bars for Upper Paleolithic assemblages from the southern Levant. Data from Rabinovich (2003).

Shifting to subsistence practices, our data show that both ibex and gazelles were hunted throughout the sequence, but the odds of ibex to gazelle increase significantly through time: by a factor of 1.4 between Layer 7 and Layer 5, and then more than double in Layers 4 and 3. It is difficult to say whether this shift in prey frequencies reflects depletion of local gazelle herds or more specialized, shorter term caprine hunting forays through time. Demographic data can be harnessed to support the second scenario, since the hunters targeted prime adults among caprines, by evidence of both tooth eruption and wear and epiphyseal fusion data; this suggests specialized hunting. Gazelles, however, appear to have been hunted in an unselective manner, with a proportion of females and young that would be typical of a recent herd. We therefore suggest a tentative reconstruction of shorter-term hunting forays focusing on goats as the environment slid into dry glacial conditions. This hypothesis is also congruent with the high frequency of weathered bones in the upper Layers 4-2, which suggests lower rates of anthropogenic deposition and, by proxy, briefer and more ephemeral occupations. It is notable that the trend of specialization in caprines contradicts the rising frequency of gazelles observed in Manot cave through time.

Carcass transport was sometimes selective and tended to bring head and upper limb portions to camp. This was not a stereotyped behavioral choice, because both feet and axis elements are present in the assemblage. In our opinion, this selective transport of rather small ungulates who, when eviscerated, weigh 10-20 kg, indicates that often carcasses were carried to the site from a distance or across rough terrain. The field butchery of smaller ungulates is different from the pattern observed in the roughly contemporary assemblage from Manot Cave, where carcasses were brought to the site complete (Yeshurun et al. 2021), and so they appear to be also in the Aurignacian of Hayonim Cave (Stiner 2005). The difference may stem from lower local carrying capacity and lower game density, which entailed longer foraging radius; or, alternatively, by the difficulty imposed by the sharp topography of the region. Carcass processing involves extensive and intensive breakage, which suggests the extraction of within bone nutrients, although this does not show in standard taphonomic comparisons of fragmentation with marrow and grease indices. A specific form of longitudinal phalanx splitting was practiced. This longitudinal is difficult to interpret as indicating resource stress (Jin & Mills, 2011) and seems to represent an idiosyncratic (for now) butchery behavior of some local Upper Paleolithic groups.

The animal remains from the site suggest a unique dual focus on different medium-sized bovids, ibex and gazelle, which are extant in the region. Caprine hunting focused on prime adults, and increased in in time; that of gazelles appears to have been random with respect to age and sex, and declines through time, a trend that we interpret as reflecting shorter stays focused on ibex hunting as the climate deteriorated. In comparison with other Final Pleistocene desert sites, NR2 is different from Epipaleolithic Tor Hamar (Klein, 1995), which is dominated by gazelles; It appears to be most similar to the Ein Aqev (D-31) assemblage described by Tchernov (1976), in which the odds of goats to gazelles are nearly even, and to which it is also similar in terms of flint industry along the lower part of the sequence. The later layers of NR2, however, with their emphasis on caprines, may be somewhat closer to the desert hunting site at LGM Wadi Madamagh (Sadhir et al. 2020), in which caprines overwhelmingly dominate the taxonomic profile.

The faunal assemblage from NR2 represents the first fair-sized assemblage from the Upper Paleolithic of the Judean Desert and the Negev that has been collected by sieving and recorded using a modern analytical protocol, and the only one associated with the Arkov-Divshon culture: The results, which suggest long ranged foraging and increased specialization in caprines through time, supply another major reference point for Upper Paleolithic human hunting adaptations in the arid regions of the southern Levant.

## Supporting information

S1 database

S2 measurements

S3 skeletal element abundance

S4 comparanda (PAST .dat file)

## Supplementary Information

Supplement 1: The archaeozoological database.

Supplement 2: Osteometric data.

Supplement 3: Skeletal element representation data.

Supplement 4: Comparative database of Levantine UP sites.

## Data, script, and code availability

Supplementary data are available in 10.5281/zenodo.6697732

Software (SEAcalc package) available in github (https://github.com/nmar79/SEAcalc.git) and at https://doi.org/10.5281/zenodo.6697892

## Acknowledgements

We are grateful for the help of our students and friends who joined us in the excavation. We also wish to thank Simon J.M. Davis and Maayan Lev for their help with taxonomic identifications. Finally, we would like to acknowledge the help of Dudi Greenberg, Jamil al-Atrash, and Noa Gordon of the National Parks Authority for facilitating this project. Version 4 of this preprint has been peer-reviewed and recommended by Peer Community In Archaeology (https://doi.org/10.24072/pci.archaeo.100017)

